# Multi-omics analysis for potential inflammation-related genes involved in tumor immune evasion via extended application of epigenetic data

**DOI:** 10.1101/2021.11.04.467249

**Authors:** Chenshen Huang, Ning Wang, Na Zhang, Zhizhan Ni, Xiaohong Liu, Hao Xiong, Huahao Xie, Boxu Lin, Bujun Ge, Qi Huang, Bing Du

## Abstract

**Background:** Accumulating evidence suggests that inflammation-related genes may play key roles in tumor immune evasion. Programmed cell death ligand 1 (PD-L1) is an important immune checkpoint involved in mediating antitumor immunity. We performed multi-omics analysis to explore key inflammation-related genes affecting the transcriptional regulation of PD-L1 expression.

**Methods:** The open chromatin region of the PD-L1 promoter was mapped using the assay for transposase-accessible chromatin using sequencing (ATAC-seq) profiles. Correlation analysis of epigenetic data (ATAC-seq) and transcriptome data (RNA-seq) were performed to identify inflammation-related transcription factors whose expression levels were correlated with the chromatin accessibility of the PD-L1 promoter. Chromatin immunoprecipitation sequencing (ChIP-seq) profiles were used to confirm the physical binding of the TF STAT2 and the predicted binding regions. We also confirmed the results of the bioinformatics analysis with cell experiments.

**Results:** We identified chr9:5449463-5449962 and chr9:5450250-5450749 as reproducible open chromatin regions in the PD-L1 promoter. Moreover, we observed a correlation between STAT2 expression and the accessibility of the aforementioned regions. Furthermore, we confirmed its physical binding through ChIP-seq profiles and demonstrated the regulation of PD-L1 by STAT2 overexpression *in vitro*. Multiple databases were also used for the validation of the results.

**Conclusion:** Our study identified STAT2 as a direct upstream TF regulating PD-L1 expression. The interaction of STAT2 and PD-L1 might be associated with tumor immune evasion in cancers, suggesting the potential value for tumor treatment.

## 1 Introduction

Immune evasion is an essential mechanism for cancer cells to circumvent immune-system mediated destruction and acquire resistance to treatment. Both laboratory and clinical studies have revealed that PD-L1 plays a key role in immune evasion. PD-L1, also known as CD274, is a co-inhibitory receptor expressed on the surface of multiple cell types, including cancer cells (Doroshow et al., 2021)(Egen et al., 2020). It can bind to programmed death-1 (PD-1) and inhibit anti-tumor immune reactions, enabling cancer cells to escape immunosurveillance.

Based on the importance of the PD-1/PD-L1 axis in immune evasion, many studies have demonstrated the remarkable clinical efficacy of anti-PD-1/PD-L1 therapy (He and Xu, 2020). However, the clinical effects of these treatments are less efficient for certain tumor types, such as non-microsatellite instability (non-MSI) colorectal cancer (Overman et al., 2017). To date, the level of PD-L1 expression in cancer cells is regarded as one of the most important factors for determining the effects of immune checkpoint therapy. Therefore, an improved understanding of PD-L1 regulation in cancer cells might be helpful for clinical cancer treatment.

Accumulating evidence has demonstrated the upregulation of PD-L1 expression during cancer pathogenesis. Inflammatory signaling is regarded as a primary mechanism involved in this complex regulatory network (Sun et al., 2018). For instance, certain pro-inflammatory factors, including type I and type II interferons, induce PD-L1 expression efficiently. Furthermore, the expression of PD-L1 could be regulated via multiple inflammation-related transcription factors (TFs), such as IRF1, STAT1, STAT3, and NF-κB (Antonangeli et al., 2020). Given that cancer-related inflammation is observed in a substantial proportion of patients, a better understanding of the relationship between inflammation and PD-L1 is currently required. Furthermore, a search for novel inflammatory TFs that regulate PD-L1 expression is warranted.

Assay for transposase-accessible chromatin using sequencing (ATAC-seq) approach uses hyperactive Tn5 transposase to comprehensively recognize chromatin accessibility at the genome level and could map open chromatin regions in gene promoters, reflecting the possibilities of TF binding (Corces et al., 2018; Nordstrom et al., 2019). Although ATAC-seq analysis only indicates the necessity of TF binding, its results could be further confirmed through other experiments.

In this study, we aimed to perform a multi-omics analysis to screen novel inflammation-related TFs involved in PD-L1 regulation, followed by laboratory verification studies. Besides transcriptome data (RNA-seq), we also aimed to analyze epigenetic data (ATAC-seq) to explore the binding of TFs. We hypothesized that STAT2 (signal transducer and activator of transcription 2) could directly bind at open chromatin regions of the PD-L1 promoter and regulate PD-L1 expression in cancer cells. The predicted binding was then validated physically through ChIP assay, and its influence in translational regulation was further confirmed in cell experiments.

## 2 Materials and methods

### 2.1 Data collection

The Cancer Genome Atlas (TCGA) datasets were accessed through the UCSC Xena database (https://xenabrowser.net/). Htseq-count profiles of 514 colon adenocarcinoma (COAD) samples were retrieved, and the corresponding clinical demographic information was also acquired. We also downloaded the Fragments Per Kilobase per Million mapped read (FPKM) profiles of the aforementioned patients with COAD. Publicly available ATAC-seq profiles were obtained from the NCI Genomic Data Commons (https://gdc.cancer.gov/about-data/publications/ATACseq-AWG). The ChIP-seq profiles were acquired from the Cistrome database (http://cistrome.org/) (Liu et al., 2011). RNA-seq profiles were obtained from the Gene Expression Omnibus (GEO, GSE50588, https://www.ncbi.nlm.nih.gov/geo/) (Cusanovich et al., 2014). For quality control, the RNA-seq results of STAT2-siRNA experiments were considered eligible only when the expression of STAT2 had a more than 4-fold difference between control group and STAT2-knockdown group.

### 2.2 Identification of inflammation-related TFs

We first downloaded the list of 1639 transcription factors (TF) from the Human Transcription Factors database (http://humantfs.ccbr.utoronto.ca/) (Lambert et al., 2018). Then, via literature search, we identified three TF families that were crucial in mediating inflammation, including nuclear factor-kB (NF-kB), interferon regulatory factors (IRFs), and signal transducers and activators of transcription (STATs) (Yu et al., 2009; Platanitis and Decker, 2018; Ni et al., 2021). Accordingly, a total of 21 TFs were identified as inflammatory TFs: NF-kB 1-2, RelB, c-Rel, IRF 1-9, STATs 1-4, 5a, 5b, and 6.

### 2.3 Chromatin accessibility analysis of PD-L1

To identify the open chromatin regions of PD-L1, peaks were visualized using the R package karyoploteR (Gel and Serra, 2017) and ChIPseeker (Yu et al., 2015), and were annotated using TxDb.Hsapiens.UCSC.Hg38. knownGene. The details of the aforementioned methods have been described in our former study (Huang et al., 2020; Ni et al., 2021).

### 2.4 Identification of potential TFs involved in PD-L1 regulation

In order to analyze transcriptional regulation of PD-L1, we used the workflow reported by Huang et al (Huang et al., 2020). Briefly, gene expression data of TFs were retrieved from the TCGA datasets. Then, we performed correlation analysis between the TF expression and the chromatin accessibility of the PD-L1 promoter region. The TFs with a p-value < 0.05 were further filtered using the Cistrome database and the GEO database.

### 2.5 Cell culture

The human colon cancer cell line DLD-1 and cervical carcinoma cell line HeLa were obtained from Shanghai Key Laboratory of Regulatory Biology, East China Normal University, Shanghai, China. Both cell lines were cultured in DMEM supplemented with 1% streptomycin–penicillin and 10% fetal bovine serum.

### 2.6 Plasmids and transfection

PcDNA3.1-STAT2 plasmid was purchased from Youbio Biological Technology Co., Ltd. (China). The transfections were performed via the calcium phosphate-DNA coprecipitation method for both DLD-1 and HeLa cells, as described previously. Equal amounts of empty vectors were transfected in the negative control group.

### 2.7 Real-time qPCR

Total RNA was extracted using the TRIzol reagent (Takara). The PrimeScript RT Master Mix Kit (Takara) was used for cDNA generation. Then, real-time qPCR was performed using SYBR Green PCR Master Mix (Yeasen). The primer sequences for each gene are listed in Supplemental Table S1.

### 2.8 Multi-database validation

To minimize the bias in bioinformatics analysis, multiple databases were used for validation, including the Timer database (http://timer.comp-genomics.org/) (Li et al., 2020), Human Protein Atlas database (https://www.proteinatlas.org/) (Uhlen et al., 2015; Uhlen et al., 2017), LinkedOmics database (https://linkedomics.org/) (Vasaikar et al., 2018), and GEPIA database (http://gepia.cancer-pku.cn/) (Tang et al., 2017).

### 2.9 Statistical analysis

P-values < 0.05 were regarded as statistically significant. Pearson and Spearman analysis was used to calculate correlation coefficients. All statistical analyses were conducted using the R software (version 3.5.1; www.r-project.org).

## 3 Results

### 3.1 Identification of open chromatin regions of the PD-L1 promoter

The overview of our study is presented in **Figure 1**. The chromatin accessibility landscape of patients with COAD was gauged from ATAC-profiles using the workflow reported by Huang et al (Huang et al., 2020). The open chromatin regions were widely expressed across the genome (**Figure 2A**). The upset plot and the vennpie plot indicated that a considerable fraction of open chromatin regions was found in gene promoters (**Figure 2B**). Furthermore, we identified chr9:5449463-5449962 (Region 1) and chr9:5450250-5450749 (Region 2) as reproducible open chromatin regions in the PD-L1 promoter across 41 patients with COAD.

**Figure 1.**
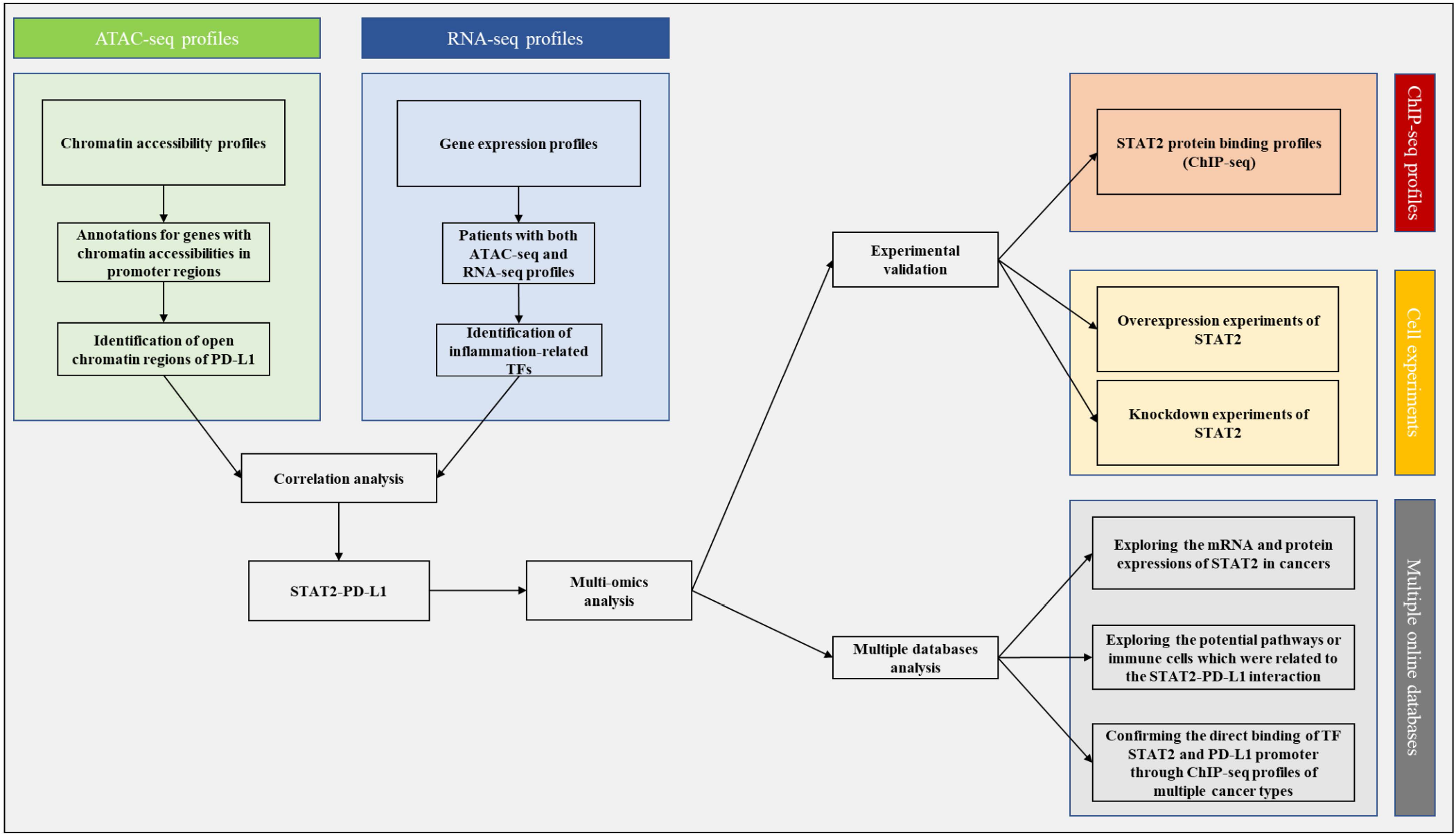
An overview of the study design.

**Figure 2.**
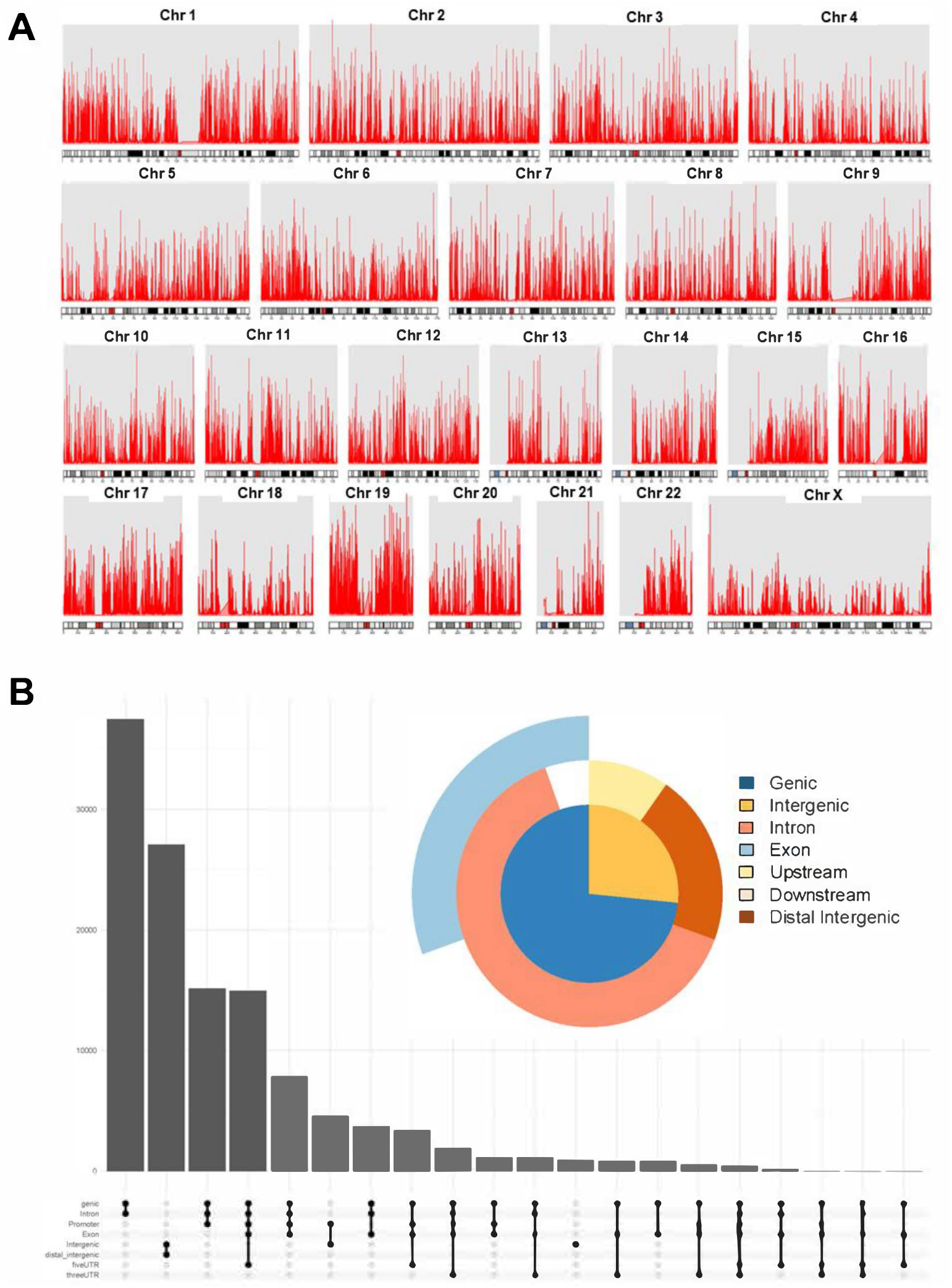
Identification of chromatin accessibilities using ATAC-seq profiles. (A) For the analysis of chromatin accessibilities, peak calling function was performed, and peaks were visualizated over whole genome. (B) The upset plot and vennpie plot revealed that a considerable fraction of open chromatin regions were around gene promoters.

### 3.2 Identification of STAT2 as an upstream factor of PD-L1 by integrative analysis

In order to explore the upstream factors of PD-L1, we combined transcriptome (RNA-seq) and epigenetics profiles (ATAC-seq) for co-analysis. Based on the list of TFs associated with inflammation, we obtained the mRNA expression of 21 TFs from RNA-seq profiles in the TCGA database (**Figure 3A**). These TFs were then filtered by correlation analysis with chromatin accessibilities of Region 1 and Region 2 (Figure 3B). After filtration, STAT2 demonstrated a remarkable correlation with both Region 1 and Region 2 (**Figure 3B, 3C**).

**Figure 3.**
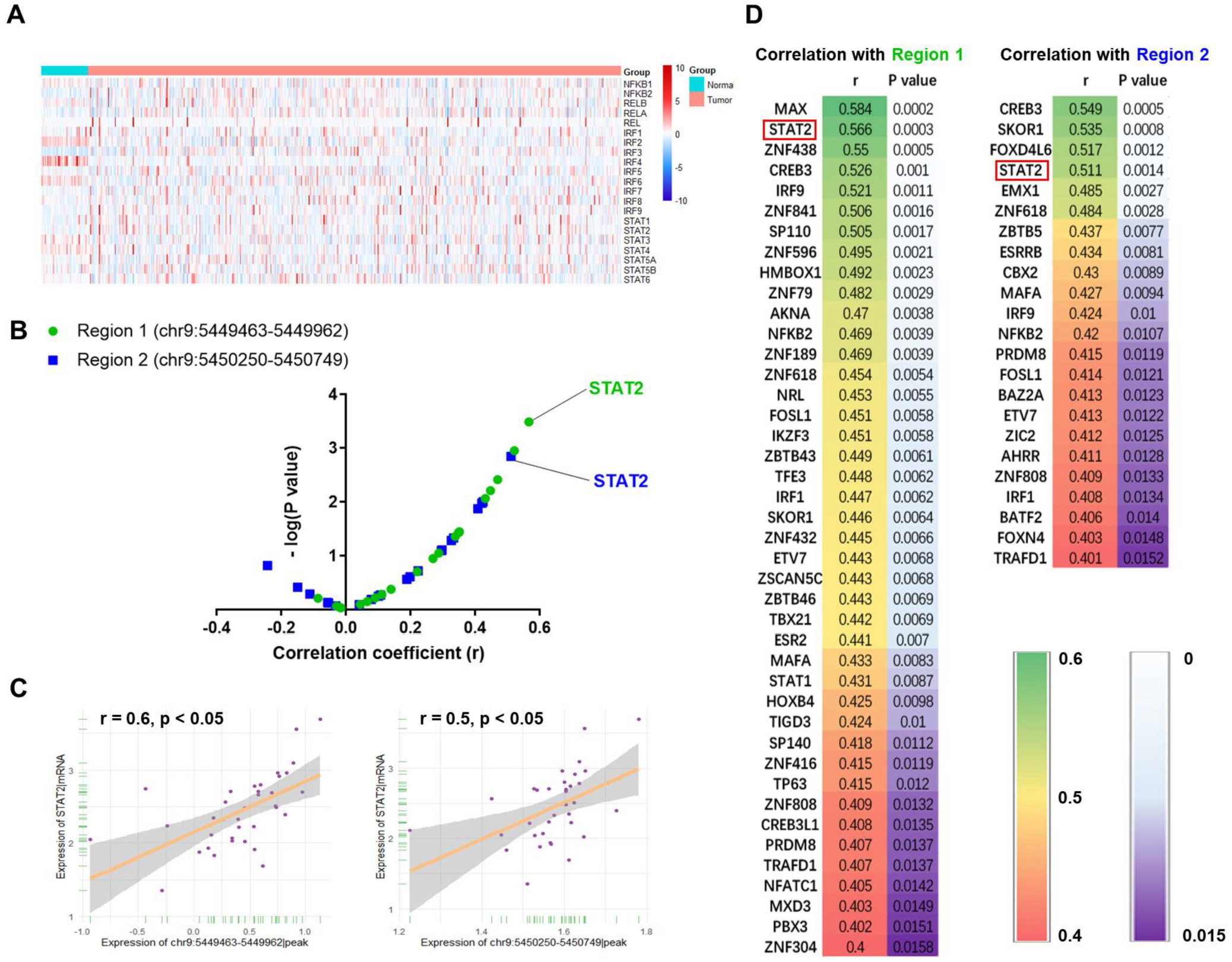
Integrative analysis of RNA-seq and ATAC-seq profiles identified STAT2 as a potential upstream for PD-L1. (A) Heatmap for gene expression of the identified 21 inflammation-related TFs, which were from NF-kB, IRFs or STATs families. The mRNA expression of all the 21 TFs could be detected in colon cancer tissues. (B) Correlation analysis revealed that STAT2 had a stronger association with chromatin accessibilities of PD-L1 promoter. (C) The STAT2 expression was significantly correlated with the chromatin accessibility of Region 1 (r = 0.6, p < 0.05). The STAT2 expression was significantly correlated with the chromatin accessibility of Region 2 (r = 0.5, p < 0.05). (D) Correlation analysis between open chromatin regions and all the database-recorded TFs was performed. The TFs, which were correlated with Region 1 or Region 2, were displayed in the heatmap (r > 0.4, p < 0.05). Compared with other TFs, STAT2 was closely related with both Region 1 (left, r = 0.6, p < 0.05) and Region 2 (right, r = 0.5, p < 0.05).

Additionally, of all database-recorded 1639 TFs (**Supplemental Table S2**), STAT2 was only secondary to TF MAX with respect to correlation with Region 1 (**Figure 3D**, left). Similarly, STAT2 also had a significant correlation with Region2 (**Figure 3D**, right). Collectively, STAT2 showed a significant correlation with the open chromatin regions of PD-L1 promoter; hence, it may physically bind to the PD-L1 promoter and regulate PD-L1 expression.

### 3.3 Validation of the physical interactions between STAT2 and PD-L1 promoter

To confirm the physical binding, we obtained STAT2 ChIP-seq profiles of colon cancer cells for validation. **Figure 4A** shows that STAT2 protein could specifically bind at Region 1 (highlighted in dark blue) and Region 2 (highlighted in light blue), confirming the physical interaction between STAT2 and PD-L1 promoter. Also, the mRNA expression of PD-L1 was significantly correlated with STAT2 expression (r = 0.53, p < 0.05, **Figure 4B**), which suggested that STAT2 might have a close relationship with PD-L1 expression. Furthermore, we obtained the RNA-seq data of cells with STAT2 knockdown from the GEO datasets. PD-L1 expression was significantly downregulated in the STAT2-knockdown group (**Figure 4C**). The analysis of the protein-protein interaction network also supported the correlation observed between STAT2 and PD-L1 (**Figure S1A**). Collectively, the integrated analysis indicated that the TF STAT2 could physically bind at the open chromatin regions of the PD-L1 promoter and might regulate PD-L1 expression.

**Figure 4.**
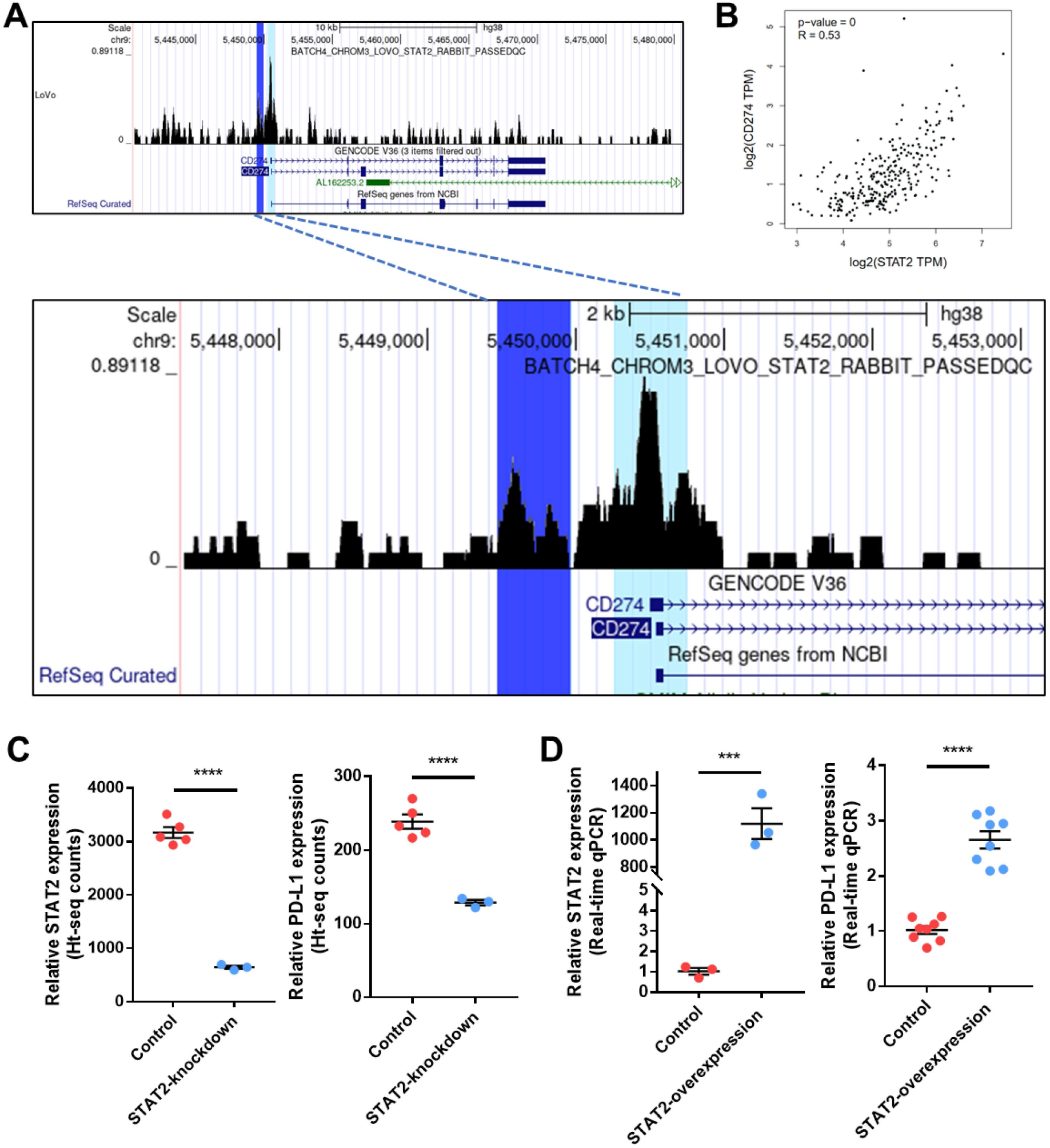
Validation of the direct regulation of STAT2 on PD-L1 through ChIP-seq profiles and cell experiments. (A) ChIP-seq profiles of STAT2 in colon cancer cell line revealed that STAT2 could directly bind to PD-L1 promoter. And there was a strong overlap between STAT2 binding sites and the predicted regions (Region 1 in dark blue, or Region 2 in light blue). (B) The mRNA expressions of STAT2 and PD-L1 were significantly correlated (r = 0.5, p < 0.05). (C) The knockdown of STAT2 could lead to down-regulation of PD-L1 significantly (p < 0.05). (D) The overexpression of STAT2 could lead to up-regulation of PD-L1 significantly (p < 0.05).

### 3.4 Validation of the results of bioinformatics analysis in colon cancer cells

To minimize the bias, we also confirmed the results of the aforementioned bioinformatics analysis with colon cancer cell line DLD-1. As expected, the transfection and overexpression of STAT2 resulted in significant upregulation of PD-L1 expression (**Figure 4D**). Similar results were also observed with HeLa cells (**Figure S1B**). Thus, via in vivo experiments, we validated that overexpression of STAT2 could upregulate PD-L1 expression in cancer cells, which was in line with the results of the bioinformatics analysis.

### 3.5 Identification of significant pathways and immune cells associated with STAT2 and PD-L1

Considering that STAT2 could be a direct upstream factor of PD-L1, we further explored associated pathways in COAD. We used the gene set variation analysis algorithm (Hanzelmann et al., 2013) to identify the expression level of genes enriched in the GO and KEGG pathway analysis. Correlation analysis was applied to explore significant pathways that were correlated with both STAT2 and PD-L1 expression (**Figure 5A, 5C**). We found that “KEGG_antigen_processing_and presentation” and “GOBP_cellular_response_to_interferon_alpha” pathways were most significantly correlated (**Figure 5B, 5D**). Therefore, we hypothesized that the interaction of STAT2 and PD-L1 might influence colon cancers in an immune-related way.

**Figure 5.**
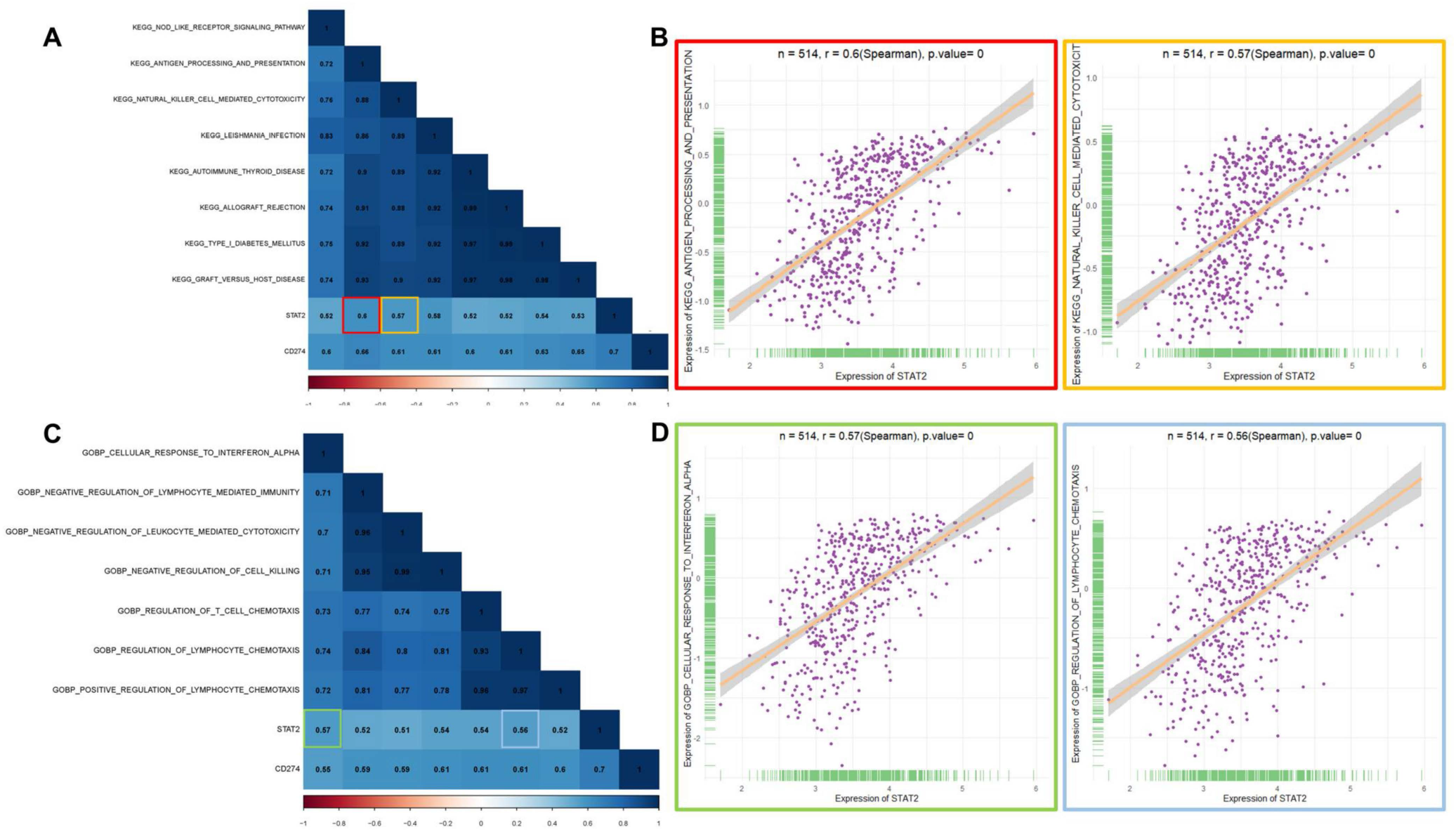
Exploring the potential pathways which were related with the interaction of STAT2 and PD-L1. (A) Correlation heatmap of STAT2, PD-L1, and the significant Kegg pathways. The pathways, which had a strong correlation with both STAT2 and PD-L1 (r > 0.4, p < 0.05), were displayed. (B) The dot plots showed the correlation between STAT2 expression, and the KEGG_antigen_processing_and_presentation pathway (marked in red, r = 0.6, p < 0.05) or KEGG_natural_killer_cell_mediated_cytotoxicit pathway (marked in yellow, r = 0.57, p < 0.05). (C) Correlation heatmap of STAT2, PD-L1, and the significant GO pathways. The pathways, which had a strong correlation with both STAT2 and PD-L1 (r > 0.4, p < 0.05), were displayed. (D) The dot plots showed the correlation between STAT2 expression, and the GOBP_cellular_response_to_interferon_alpha pathway (marked in green, r = 0.57, p < 0.05) or GOBP_regulation_of_lymphocyte_chemotaxis pathway (marked in blue, r = 0.56, p < 0.05).

Considering this hypothesis, we analyzed tumor-infiltrating immune cells in colon cancers. With the clustering function of R package corrplot, we found that macrophages might be potentially associated with both STAT2 and PD-L1 (**Figure 6A**). Furthermore, besides clustering function, we also explored the estimated immune cells using correlation analyses. And we found that algorithm-estimated macrophages and T cells were statistically correlated with both STAT2 and PD-L1 (r > 0.4, p < 0.05, **Figure 6B**). Thus, based on the analysis above, macrophages might play a role in the interaction between STAT2 and PD-L1. However, limited to bioinformatics methods, we only observed a correlation among macrophages, STAT2, and PD-L1. The underlying mechanisms still required further elucidation through laboratory experiments.

**Figure 6.**
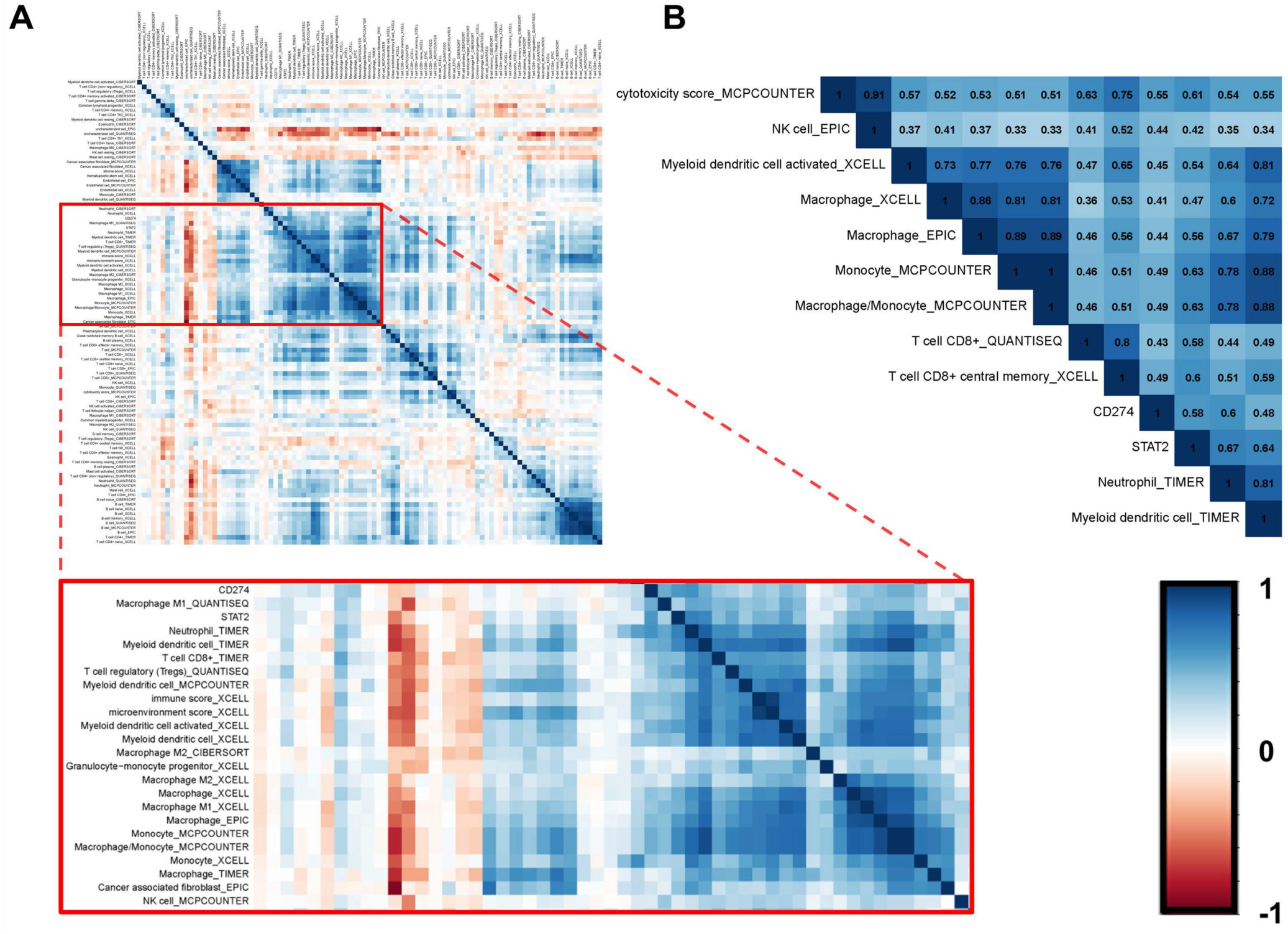
Exploring the potential immune cells which were related with the interaction of STAT2 and PD-L1. (A) Correlation heatmap of STAT2, PD-L1, and the significant tumor infiltrating immune cells. After the clustering through R package corrplot, STAT2 and PD-L1 were found to be potentially associated with multiple cell types, especially macrophages. (B) The correlation heatmap showed the immune cells that had a strong correlation with both STAT2 and PD-L1 (r > 0.4, p < 0.05).

### 3.6 Multi-database validation

To confirm the bioinformatics results, we used multiple databases for validation. Profiles from Xena database showed that STAT2 was widely expressed in multiple cancer types, including colon cancers (**Figure 7A**). Moreover, the HPA database indicated that in the tumor microenvironment of colon cancers, STAT2 could be detected in multiple cell types, including cancer cells. Although there is no significant difference of STAT2 protein expression between colon cancer tissues and normal tissues, we found that the expression level of STAT2 in normal endothelial cells was relatively lower (**Figure 7B**). In addition, the Linkedomics database demonstrated that the STAT2 expression was significantly lower in patients with MSS (non-MSI) colon cancer compared to that in patients with MSI-H colon cancer (**Figure 7C**). To confirm the direct binding of TF STAT2 and PD-L1 promoter, we also obtained STAT2 ChIP-seq profiles of multiple types of cancer cell lines, including GM12878, K562, and LoVo. The TF STAT2 were confirmed to directly bind to the predicted regions of PD-L1 promoter (**Figure 7D**).

**Figure 7.**
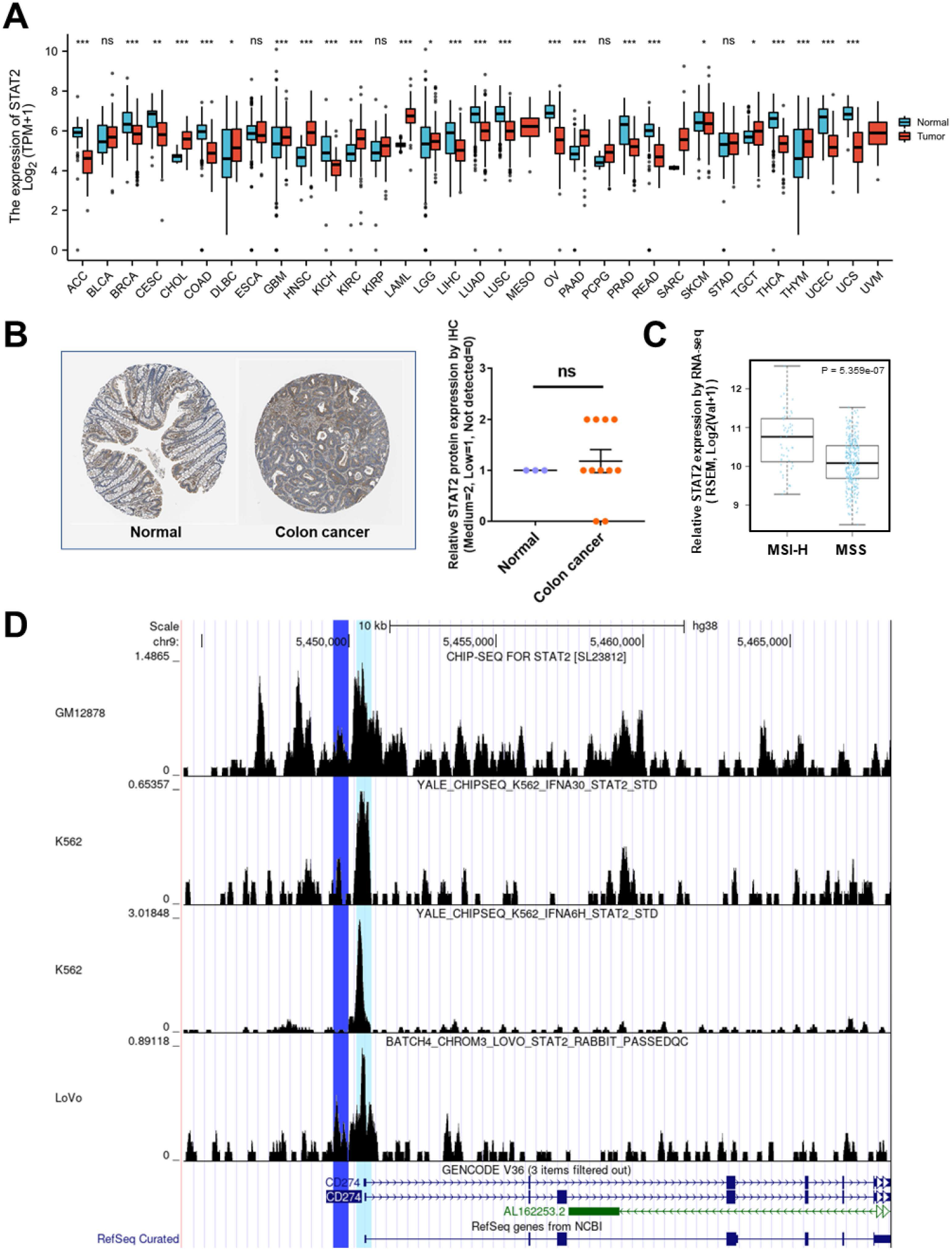
Multiple databases were used for validation. (A) Box plots of STAT2 expression in different cancers from the TCGA and GTEx databases accessed by Xena. STAT2 was widely expressed across multiple cancer types (B) Immunohistochemical results of the protein expression of STAT2 in patients with colon cancers via the HPA database. (C) Box plots revealed a significant difference of STAT2 expression between MSI-H and MSS (no-MSI) colon cancer (p < 0.05). (D) The overlap of STAT2 binding sites and the predicted region (Region 1 in dark blue, or Region 2 in light blue) was validated in STAT2 ChIP-seq profiles of multipe cell lines, including GM12878, K562, and LoVo.

## 4 Discussion

With the understanding of immune evasion mechanisms, immune checkpoint therapy has been developed as an important clinical strategy for cancer treatment. Among various such strategies, anti-PD-1/PD-L1 therapy is one of the most extensively examined strategies. Considering that the level of PD-L1 expression in cancer cells is closely associated with clinical efficacy, there is an urgent need to elucidate mechanisms underlying PD-L1 regulation. Cancer-induced chronic inflammation is very common among patients and affects PD-L1 expression via multiple pathways, including transcriptional regulation. However, to date, its regulatory mechanism has not been fully clarified. Therefore, we aimed to screen undiscovered inflammatory TFs that were direct upstream factors of PD-L1 and to further confirm the results via laboratory validation.

Based on the integrated analysis of transcriptome and epigenetic profiles, we proposed that the TF STAT2 could directly bind at the PD-L1 promoter. We also identified the potential binding sites chr9:5449463-5449962 and chr9:5450250-5450749. Upon analyzing ChIP-seq profiles of STAT2, we confirmed the physical binding within the predicted region. Subsequently, we further verified our results at the cellular level. We demonstrated that the overexpression of STAT2 can significantly upregulate PD-L1 expression in cancer cell lines DLD-1 and HeLa. Based on the GEO database, we found that STAT2-siRNA could significantly inhibit PD-L1 expression. Taken together, based on the results of this study, we propose that STAT2 is a direct upstream factor of PD-L1.

STAT2 is known for its role in immunomodulatory reactions and anti-viral immunity. It is significantly different from other members of the STAT family. For instance, although other members can be activated by multiple cytokines, including type I and II interferons, STAT2 is primarily activated by type I interferon. Moreover, it is involved in mediating inflammatory pathways and acts as a co-factor (Stark and Darnell, 2012). Thus, the regulation of STAT2 might not have a significant impact on anti-tumor immunity. Therefore, if PD-L1 expression is regulated via STAT2, the combination therapy targeting STAT2 might be more effective.

In this study, we demonstrated the regulation of PD-L1 expression mediated by STAT2 via multi-omics analysis and laboratory validation. Over past decades, the regulation of PD-L1 expression by various inflammatory TFs has been reported; however, our study is the first to report that the TF STAT2 could directly regulate PD-L1 expression. We also confirmed the physical binding of the TF STAT2 and PD-L1 promoter based on ChIP-seq results. Interestingly, a study performed by Angel Garcia-Diaz et al. (Garcia-Diaz et al., 2017) used mutagenesis to delete the predicted binding sites of STAT2 in a firefly luciferase reporter plasmid comprising the PD-L1 promoter. The results demonstrated that interferon-induced luciferase expression was remarkably decreased in the transfected cells. To some extent, the results of this experiment supplemented the findings of our study, which provided strong evidence for the direct binding of the TF.

In addition, Angel Garcia-Diaz et al. (Garcia-Diaz et al., 2017) not only found that the deletion of STAT2 putative binding site could affect luciferase expression, but also observed similar results upon deletion of IRF1 putative site. Indicating the importance of the JAK1/JAK2-STAT1/STAT2/STAT3-IRF1 axis, they revealed that IRF1 was the potential TF that regulated PD-L1 directly. In our study, we focused on the direct regulation of PD-L1 expression by STAT2, providing an improved understanding of the JAK1/JAK2-STAT1/STAT2/STAT3-IRF1 axis. As a direct upstream factor of both PD-L1 and IRF1, STAT2 showed a great promise for anti-PD-1/PD-L1 immunotherapy.

Despite the increasing number of studies examining STAT2, the effects of STAT2 in anti-tumor immunity remain controversial (Verhoeven et al., 2020). For instance, Yue et al. (Yue et al., 2015) demonstrated the acceleration of tumor growth in mice with STAT2 knockout. Wang et al. (Wang et al., 2003) confirmed the anti-tumor activity of STAT2 in a mouse model. Gamero et al. (Gamero et al., 2010) used models of inflammation-induced cancers to demonstrate that STAT2 might promote colorectal and skin carcinogenesis. Considering the controversial roles of STAT2, our study might provide further clarifications with respect to the role of STAT2 in cancer. We found that since STAT2 can upregulate PD-L1 expression on the surface of cancer cells, it can aid the cancer cells in escaping immunosurveillance.

Currently, the anti-PD-1/PD-L1 treatment demonstrates low efficiency for patients with non-MSI colon cancers (Overman et al., 2017; Sun et al., 2018). Based on the LinkedOmics database, we found that the STAT2 expression was significantly lower in no-MSI colon cancers. Thus, we hypothesized that the combination treatment targeting STAT2 might increase the efficacy of anti-PD-1/PD-L1 treatment in patients diagnosed with this cancer subtype.

Certain limitations of this study need to be addressed. First, most of our bioinformatics results were based on the data on colon cancers, which were used to perform integrated analysis. On the one hand, the multi-omics analysis would be more reliable when using profiles from the same cancer type. On the other hand, these results might be specific to colon cancers. To minimize the bias, we used multiple databases to validate our results at the pan-cancer level, and we also used other types of cell lines in subsequent experiments. Second, owing to experimental limitations, we could not reproduce the luciferase experiment performed by Angel Garcia-Diaz et al., which may support our conclusions.

Despite the aforementioned limitations, our study was the first to highlight the direct regulation of PD-L1 expression mediated by the TF STAT2. We performed both bioinformatics and laboratory analysis to validate our results. Future studies should further validate the interaction of STAT2 and PD-L1 with larger data sizes, different cancer cell lines, and the STAT2 knockdown mouse model. The potential therapeutic value of the combination treatment should also be analyzed further.

### Conclusions

Our study identified STAT2 as a direct upstream TF that regulates PD-L1 expression, suggesting its potential to be used as a therapeutic target for tumor treatment.

## 6 Author Contributions

Conception/ design: CH

Collection and/or assembly of data: CH, NW, NZ, HX, BL

Data analysis and interpretation: CH, NZ, XL, HX

Manuscript writing: CH, NW, BG, QH, BD

All authors read and approved the final manuscript.

## 8

**Figure S1 |** Validation of the interaction between STAT2 and PD-L1.

(A) The analysis of protein-protein interactions, based on GeneMANIA database, indicated a potential interaction between STAT2 and PD-L1.

(B) In cancer cell line HeLa, the overexpression of STAT2 could lead to up-regulation of PD-L1 significantly (p < 0.05).

## 9 Supplementary Material

**Supplemental Table S1 |** Primer sequences used in this study.

**Supplemental Table S2 |** List of 1639 TFs from Human Transcription Factors database.

## References

Antonangeli, F., Natalini, A., Garassino, M.C., Sica, A., Santoni, A., and Di Rosa, F. (2020). Regulation of PD-L1 Expression by NF-kappaB in Cancer. Front Immunol 11, 584626. doi: 10.3389/fimmu.2020.584626.

Corces, M.R., Granja, J.M., Shams, S., Louie, B.H., Seoane, J.A., Zhou, W., et al. (2018). The chromatin accessibility landscape of primary human cancers. Science 362(6413). doi: 10.1126/science.aav1898.

Cusanovich, D.A., Pavlovic, B., Pritchard, J.K., and Gilad, Y. (2014). The functional consequences of variation in transcription factor binding. PLoS Genet 10(3), e1004226. doi: 10.1371/journal.pgen.1004226.

Doroshow, D.B., Bhalla, S., Beasley, M.B., Sholl, L.M., Kerr, K.M., Gnjatic, S., et al. (2021). PD-L1 as a biomarker of response to immune-checkpoint inhibitors. Nat Rev Clin Oncol 18(6), 345–362. doi: 10.1038/s41571-021-00473-5.

Egen, J.G., Ouyang, W., and Wu, L.C. (2020). Human Anti-tumor Immunity: Insights from Immunotherapy Clinical Trials. Immunity 52(1), 36–54. doi: 10.1016/j.immuni.2019.12.010.

Gamero, A.M., Young, M.R., Mentor-Marcel, R., Bobe, G., Scarzello, A.J., Wise, J., et al. (2010). STAT2 contributes to promotion of colorectal and skin carcinogenesis. Cancer Prev Res (Phila) 3(4), 495–504. doi: 10.1158/1940-6207.CAPR-09-0105.

Garcia-Diaz, A., Shin, D.S., Moreno, B.H., Saco, J., Escuin-Ordinas, H., Rodriguez, G.A., et al. (2017). Interferon Receptor Signaling Pathways Regulating PD-L1 and PD-L2 Expression. Cell Rep 19(6), 1189–1201. doi: 10.1016/j.celrep.2017.04.031.

Gel, B., and Serra, E. (2017). karyoploteR: an R/Bioconductor package to plot customizable genomes displaying arbitrary data. Bioinformatics 33(19), 3088–3090. doi: 10.1093/bioinformatics/btx346.

Hanzelmann, S., Castelo, R., and Guinney, J. (2013). GSVA: gene set variation analysis for microarray and RNA-seq data. BMC Bioinformatics 14, 7. doi: 10.1186/1471-2105-14-7.

He, X., and Xu, C. (2020). Immune checkpoint signaling and cancer immunotherapy. Cell Res 30(8), 660–669. doi: 10.1038/s41422-020-0343-4.

Huang, C., Huang, R., Chen, H., Ni, Z., Huang, Q., Huang, Z., et al. (2020). Chromatin Accessibility Regulates Gene Expression and Correlates With Tumor-Infiltrating Immune Cells in Gastric Adenocarcinoma. Front Oncol 10, 609940. doi: 10.3389/fonc.2020.609940.

Lambert, S.A., Jolma, A., Campitelli, L.F., Das, P.K., Yin, Y., Albu, M., et al. (2018). The Human Transcription Factors. Cell 172(4), 650–665. doi: 10.1016/j.cell.2018.01.029.

Li, T., Fu, J., Zeng, Z., Cohen, D., Li, J., Chen, Q., et al. (2020). TIMER2.0 for analysis of tumor-infiltrating immune cells. Nucleic Acids Res 48(W1), W509–W514. doi: 10.1093/nar/gkaa407.

Liu, T., Ortiz, J.A., Taing, L., Meyer, C.A., Lee, B., Zhang, Y., et al. (2011). Cistrome: an integrative platform for transcriptional regulation studies. Genome Biol 12(8), R83. doi: 10.1186/gb-2011-12-8-r83.

Ni, Z., Huang, C., Zhao, H., Zhou, J., Hu, M., Chen, Q., et al. (2021). PLXNC1: A Novel Potential Immune-Related Target for Stomach Adenocarcinoma. Front Cell Dev Biol 9, 662707. doi: 10.3389/fcell.2021.662707.

Nordstrom, K.J.V., Schmidt, F., Gasparoni, N., Salhab, A., Gasparoni, G., Kattler, K., et al. (2019). Unique and assay specific features of NOMe-, ATAC- and DNase I-seq data. Nucleic Acids Res 47(20), 10580–10596. doi: 10.1093/nar/gkz799.

Overman, M.J., McDermott, R., Leach, J.L., Lonardi, S., Lenz, H.J., Morse, M.A., et al. (2017). Nivolumab in patients with metastatic DNA mismatch repair-deficient or microsatellite instability-high colorectal cancer (CheckMate 142): an open-label, multicentre, phase 2 study. Lancet Oncol 18(9), 1182–1191. doi: 10.1016/S1470-2045(17)30422-9.

Platanitis, E., and Decker, T. (2018). Regulatory Networks Involving STATs, IRFs, and NFkappaB in Inflammation. Front Immunol 9, 2542. doi: 10.3389/fimmu.2018.02542.

Stark, G.R., and Darnell, J.E. Jr., (2012). The JAK-STAT pathway at twenty. Immunity 36(4), 503–514. doi: 10.1016/j.immuni.2012.03.013.

Sun, C., Mezzadra, R., and Schumacher, T.N. (2018). Regulation and Function of the PD-L1 Checkpoint. Immunity 48(3), 434–452. doi: 10.1016/j.immuni.2018.03.014.

Tang, Z., Li, C., Kang, B., Gao, G., Li, C., and Zhang, Z. (2017). GEPIA: a web server for cancer and normal gene expression profiling and interactive analyses. Nucleic Acids Res 45(W1), W98–W102. doi: 10.1093/nar/gkx247.

Uhlen, M., Fagerberg, L., Hallstrom, B.M., Lindskog, C., Oksvold, P., Mardinoglu, A., et al. (2015). Proteomics. Tissue-based map of the human proteome. Science 347(6220), 1260419. doi: 10.1126/science.1260419.

Uhlen, M., Zhang, C., Lee, S., Sjostedt, E., Fagerberg, L., Bidkhori, G., et al. (2017). A pathology atlas of the human cancer transcriptome. Science 357(6352). doi: 10.1126/science.aan2507.

Vasaikar, S.V., Straub, P., Wang, J., and Zhang, B. (2018). LinkedOmics: analyzing multi-omics data within and across 32 cancer types. Nucleic Acids Res 46(D1), D956–D963. doi: 10.1093/nar/gkx1090.

Verhoeven, Y., Tilborghs, S., Jacobs, J., De Waele, J., Quatannens, D., Deben, C., et al. (2020). The potential and controversy of targeting STAT family members in cancer. Semin Cancer Biol 60, 41–56. doi: 10.1016/j.semcancer.2019.10.002.

Wang, J., Pham-Mitchell, N., Schindler, C., and Campbell, I.L. (2003). Dysregulated Sonic hedgehog signaling and medulloblastoma consequent to IFN-alpha-stimulated STAT2-independent production of IFN-gamma in the brain. J Clin Invest 112(4), 535–543. doi: 10.1172/JCI18637.

Yu, G., Wang, L.G., and He, Q.Y. (2015). ChIPseeker: an R/Bioconductor package for ChIP peak annotation, comparison and visualization. Bioinformatics 31(14), 2382–2383. doi: 10.1093/bioinformatics/btv145.

Yu, H., Pardoll, D., and Jove, R. (2009). STATs in cancer inflammation and immunity: a leading role for STAT3. Nat Rev Cancer 9(11), 798–809. doi: 10.1038/nrc2734.

Yue, C., Xu, J., Tan Estioko, M.D., Kotredes, K.P., Lopez-Otalora, Y., Hilliard, B.A., et al. (2015). Host STAT2/type I interferon axis controls tumor growth. Int J Cancer 136(1), 117–126. doi: 10.1002/ijc.29004.

