## Supplementary figures and images for "Multi-omics analysis for potential inflammation-related genes involved in tumor immune evasion via extended application of epigenetic data"

### Supplemental Figure S1

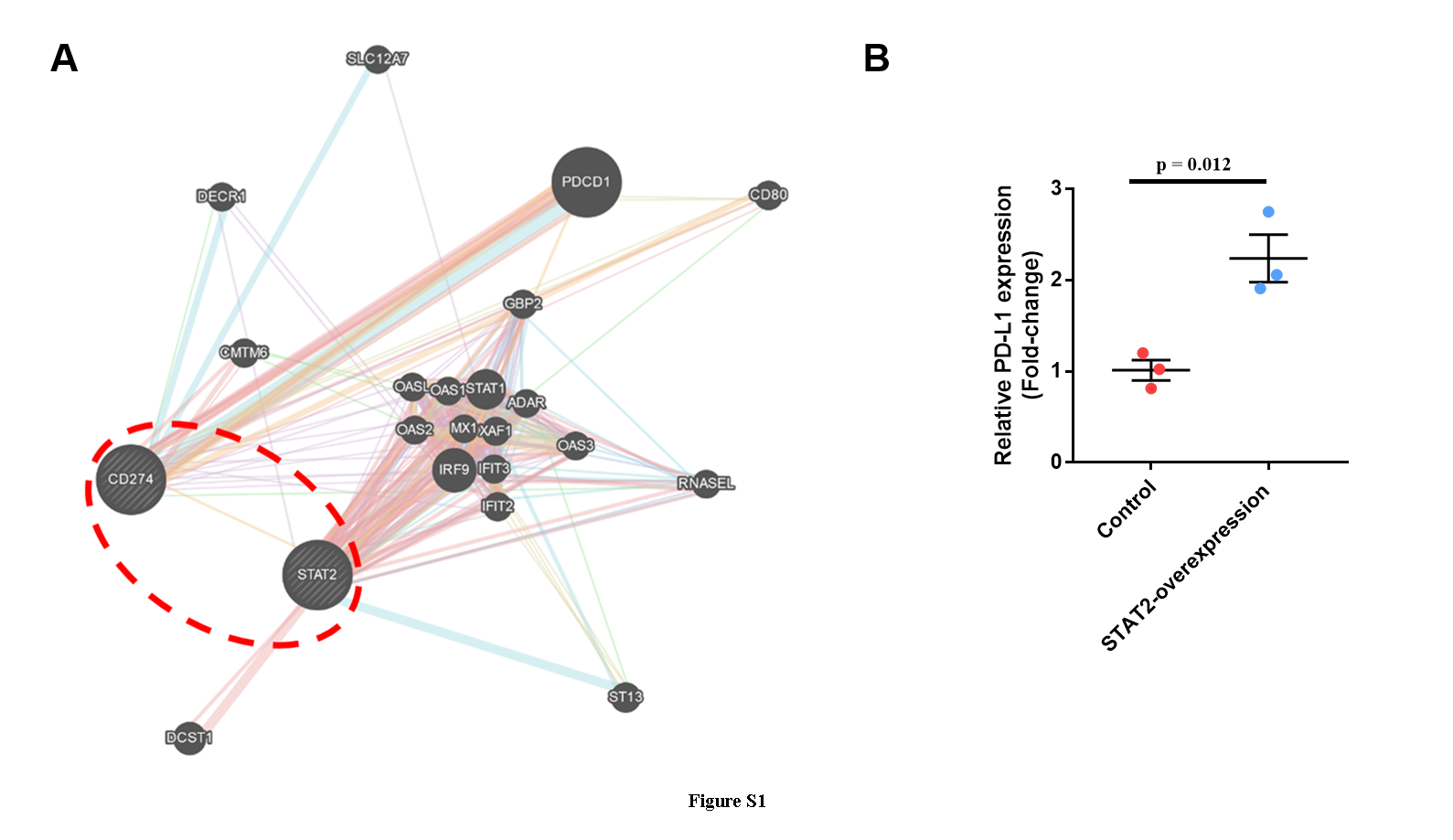
