## Supplemental Table S1 for "Multi-omics analysis for potential inflammation-related genes involved in tumor immune evasion via extended application of epigenetic data"

Supplemental Table S1. Primer sequences used in this study.

| gene | primer |
| --- | --- |
| human PD-L1 | Forward 5’-TGGCATTTGCTGAACGCATTT-3’ |
|  | Reverse 5’-TGCAGCCAGGTCTAATTGTTTT-3’ |
| human STAT2 | Forward 5’-CCAGCTTTACTCGCACAGC-3’ |
|  | Reverse 5’-AGCCTTGGAATCATCACTCCC-3’ |
| human GAPDH | Forward 5’-ACCCAGAAGACTGTGGATGG-3’ |
|  | Reverse 5’-TTCAGCTCAGGGATGACCTT-3’ |
| human β-actin | Forward 5’-ACACTGTGCCCATCTACGAG-3’ |
|  | Reverse 5’-TCAACGTCACACTTCATGATG-3’ |
